# D936Y and Other Mutations in the Fusion Core of the SARS-Cov-2 Spike Protein Heptad Repeat 1 Undermine the Post-Fusion Assembly

**DOI:** 10.1101/2020.06.08.140152

**Authors:** Luigi Cavallo, Romina Oliva

**Affiliations:** King Abdullah University of Science and Technology (KAUST), Physical Sciences and Engineering Division, Kaust Catalysis Center, Thuwal 23955-6900, Saudi Arabia; Department of Sciences and Technologies, University Parthenope of Naples, Centro Direzionale Isola C4, I-80143, Naples, Italy

**Keywords:** COVID-19, membrane fusion, mutations, HR1, helical bundle, molecular modeling, salt-bridge, aromatic, infectivity

## Abstract

The iconic “red crown” of the severe acute respiratory syndrome coronavirus 2 (SARS-CoV-2) is made of its spike (S) glycoprotein. The S protein is the Trojan horse of coronaviruses, mediating their entry into the host cells. While SARS-CoV-2 was becoming a global threat, scientists have been accumulating data on the virus at an impressive pace, both in terms of genomic sequences and of three-dimensional structures. On April 21^st^, the GISAID resource had collected 10,823 SARS-CoV-2 genomic sequences. We extracted from them all the complete S protein sequences and identified point mutations thereof. Six mutations were located on a 14-residue segment (929-943) in the “fusion core” of the heptad repeat 1 (HR1). Our modeling in the pre- and post-fusion S protein conformations revealed, for three of them, the loss of interactions stabilizing the post-fusion assembly. On May 29^th^, the SARS-CoV-2 genomic sequences in GISAID were 34,805. An analysis of the occurrences of the HR1 mutations in this updated dataset revealed a significant increase for the S929I and S939F mutations and a dramatic increase for the D936Y mutation, which was particularly widespread in Sweden and Wales/England. We notice that this is also the mutation causing the loss of a strong inter-monomer interaction, the D936-R1185 salt bridge, thus clearly weakening the post-fusion assembly.

## Introduction

Coronavirus Disease 2019 (COVID-19) is caused by the severe acute respiratory syndrome coronavirus 2 (SARS-CoV-2). SARS-CoV-2 is a novel virus belonging to the β genus coronaviruses, which also include two highly pathogenic human viruses identified in the last two decades, the severe acute respiratory syndrome coronavirus (SARS-CoV) and the Middle East respiratory syndrome coronavirus (MERS-CoV) (1–3).

Coronaviruses are named after the protruding spike (S) glycoproteins on their envelope, giving a crown (*corona* in latin) shape to the virions (4). Of the four structural proteins of coronavirues, S, envelope (E), membrane (M), and nucleocapsid (N), the S protein is the one playing a key role in mediating the viral entry into the host cells (5–7), making it one of the main targets for the development of therapeutic drugs and vaccines (8–14). Comprised of two functional subunits, S1 and S2, it first binds to a host receptor through the receptor-binding domain (RBD) in the S1 subunit and then fuses the viral and host membranes through the S2 subunit (7,15). In the prefusion conformation, the SARS-CoV-2 S protein forms homotrimers protruding from the viral surface, where its RBD binds to the angiotensin-converting enzyme 2 (ACE2) receptor on the host cell surface (1) (like the SARS-CoV homolog (16), and differently from MERS-CoV S, which recognizes a different receptor, the dipeptidyl peptidase 4 (17)). Receptor binding and proteolytic processing by cellular proteases then cause S1 to dissociate and S2 to undergo large-scale conformational changes towards a stable structure, bringing viral and cellular membranes into close proximity for fusion and infection (7,15,18).

While the outbreak of COVID-19 was rapidly spreading all over the world, affecting millions of people and becoming a global threat, laboratories worldwide promptly started to sequence a large number of SARS-CoV-2 genomes. All the available genomic data is accessible through the Global Initiative on Sharing All Influenza Data (GISAID) website, an invaluable open access resource (19,20). Simultaneously, crucial structural knowledge has been achieved on SARS-CoV-2, especially regarding the S protein. 3D structures are now available from the Protein Data Bank (PDB) (21) for the SARS-CoV-2 S protein in the pre-fusion conformation, also bound to the ACE2 receptor (22–28), and for the post-fusion core of its S2 subunit in the postfusion conformation (29).

On April 21^st^ 2020, 4 months after the first sequencing (30), 10,823 genomic sequences of SARS-CoV-2 were available from GISAID. Therefore, we considered the time ripe for an assessment of the mutational spectrum of the SARS-CoV-2 spike protein. To this aim, we extracted all the complete S protein sequences from the GISAID 21^st^ April dataset and identified all the mutations occurring in at least 2 identical sequences (see Table S1). From this analysis, a 14-amino acid segment in the fusion core of the heptad repeat 1 (HR1) emerged as a hotspot for mutations. While the mutations we identified corresponded to a 1 mutation every 12 positions along the protein sequence, as many as 6 amino acids were found to be mutated in the above 14-amino acid segment: S929, D936, L938, S939, S940 and S943.

After the proteolytic processing, in the post-fusion conformation, the S protein HR1 and HR2 motifs interact with each other to form a six-helix bundle (6-HB), which promotes initiation of the viral and cellular membranes fusion. The HR1 “fusion core” is named after its role in giving many interactions with HR2 in the post-fusion conformation, thus playing a key role in the virus infectivity (31). Based on the structural location of the above highly concentrated mutations and on their nonconservative nature, we considered them of particular interest and decided to further investigate their structural basis, both in the pre- and post-fusion conformation, as well as their sequencing dates and geographical distribution. As we show in the following, as many as three of them are responsible for the loss of inter-monomer H-bonds in the post-fusion conformation, while one of them, S943P, would introduce unexpected structural strain in the pre-fusion conformation.

A search in the GISAID resource updated to May 29^th^ showed a significant increase in occurrences especially for one mutant, D936Y, unreported to date, which has become a common variant in some European countries, especially Sweden. It is also the mutant having the most significant structural role, causing the loss of an intermonomer salt bridge in the post-fusion assembly.

## Methods

### Identification of mutations

We downloaded the 10,823 genomic sequences available from GISAID on April 21^st^ 2020. From these sequences, we extracted the nucleotide sequences of the spike protein and translated them to protein sequences with in-house scripts. Nucleotides sequences featuring an internal stop codon, having at least one undefined (“N”) nucleotide or resulting in spike proteins of length different from 1,273 amino acids were discarded. Sequences annotated as pangolin, bat or canine were also discarded. The remaining 7,692 protein sequences were further analysed. First, we clustered them in sets of identical sequences with CD-HIT (32), obtaining 120 clusters of at least 2 sequences and 245 unique sequences. As a reference system for further analyses, we used the first dated (on December 24^th^ 2019) genomic sequence in GISAID, isolated and sequenced in Wuhan (Hubei, China) (30). Then, upon alignment to the reference sequence, we identified point mutations in all the sets of at least two sequences.

We downloaded again the 34,805 genomic sequences available from GISAID on May 29^th^ 2020 (gisaid_hcov-19_2020_05_29_14) and followed the above pipeline to extract 23,332 complete 1273-residue long S protein sequences. We then recorded the presence and frequency in them of any mutation occurring in the fusion core of the HR1 (residues 929-949) with in-house scripts.

### Mutants modelling and analysis

Mutants 3D models were built using the mutate_model module of the Modeller 9v11 program (33). This is an automated method for modelling point mutations in protein structures, which includes an optimisation procedure of the mutated residue in its environment, beginning with a conjugate gradients minimisation, continuing with molecular dynamics with simulated annealing and finishing again by conjugate gradients. The used force field is CHARM-22, for details see Reference (34). Models for mutants in the pre-fusion conformation were built starting from the EM structure of the pre-fusion trimeric conformation (PDB ID: 6VSB, resolution 3.46 Å, (22)). Models for mutants in the post-fusion conformation were built starting from the X-ray structure of the S2 subunit fusion core (PDB ID: 6LXT, resolution 2.90 Å, (29)). Molecular models were analysed and visually inspected with Pymol (35). The COCOMAPS web server (36) was used to analyse the inter-chain contacts and H-bonds as well as the residues accessibility to the solvent.

## Results and Discussion

We downloaded all the SARS-CoV-2 genomic sequences from the GISAID resource on April 21^st^ 2020, extracted from them 7,692 complete S protein sequences and identified all the point mutations occurring in at least two identical sequences (see Methods). The 111 mutations we identified, occurring at 105 different positions spread all over the protein sequence, are reported in Table S1, with the relative number of occurrences.

While the mutations we identified were spaced on average 12 positions along the protein sequence, a segment of 14 amino acids harboured 6 mutations, at positions 929, 936, 938, 939, 940 and 943, proposing itself as a mutational hotspot. This sequence segment is part of the “fusion core” of the HR1, in the protein S2 subunit. The HR1 of coronaviruses S proteins undergoes one of the most notable rearrangements within the protein between the pre- and post-fusion conformations. In the post-fusion conformation, in fact, it experiences a refolding of the pre-fusion multiple helices and intervening regions into a single continuous helix (Figure 1). As already mentioned, three of these long helices then form a 6HB with three HR2 helical motifs (18,29,31). The HR1 and its “fusion core” in particular thus play a crucial role in the virus infectivity.

**Figure 1.**
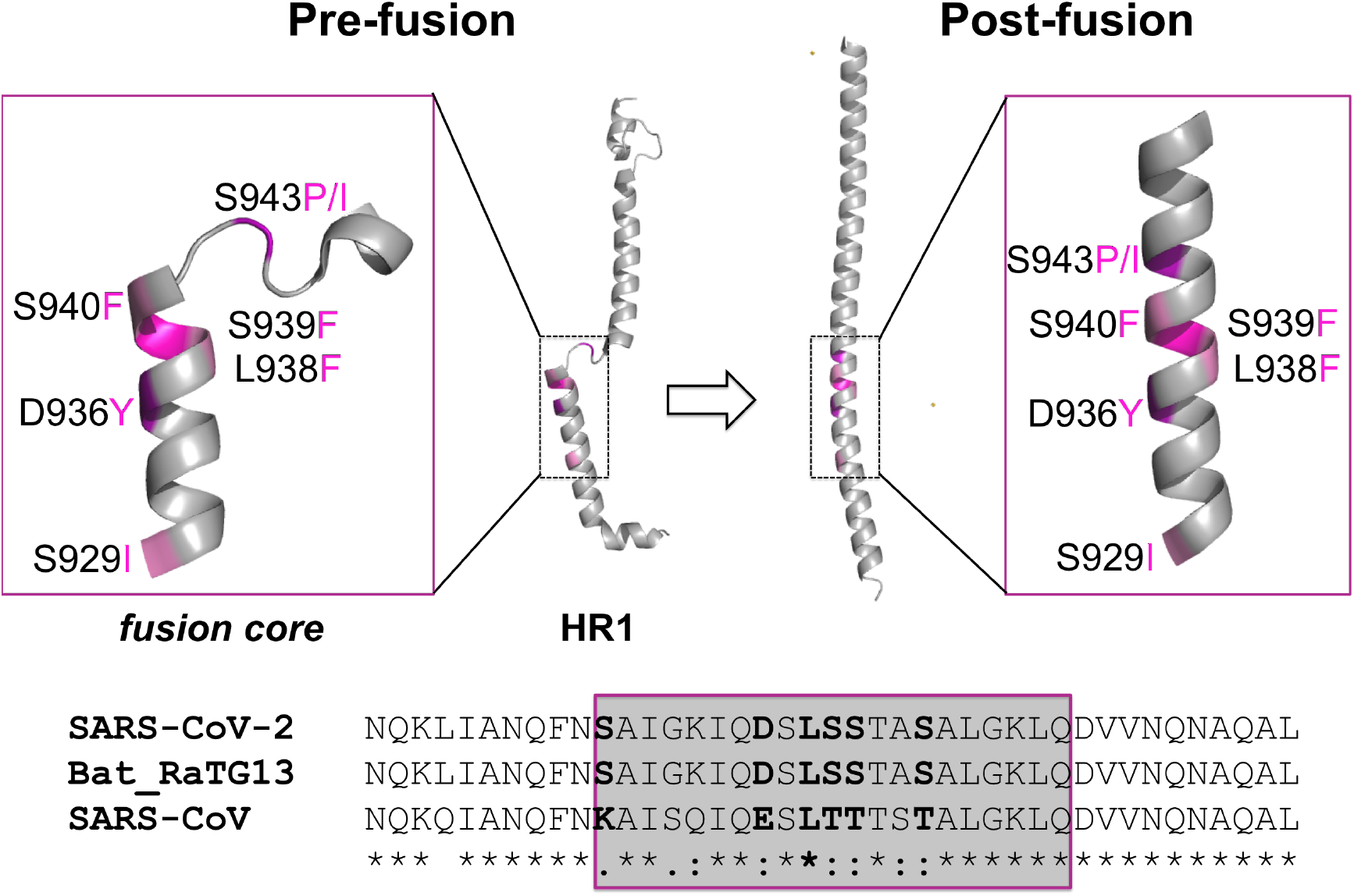
*Top:* Cartoon representation of the SARS-CoV-2 S protein HR1 and its fusion core (insets) in the pre- and post-fusion conformations (PDB IDs: 6VSB and 6LXT). Discussed mutations are colored in a purple-to-pink scale, depending on their frequency, and labelled. *Bottom:* Sequence alignment of the HR1 fusion core (framed) and 10 residues up- and down-stream in the S protein of SARS-CoV-2, bat coronavirus RaTG13 (protein_ID: QHR63300.2) and SARS-CoV (protein_ID: AAP13441.1).

### HR1 “fusion core” mutations: update on April 21^st^

The following 6 mutations were identified in the fusion core of the HR1 on April 21^st^ 2020: S929I, D936Y, L938F, S939F, S940F, S943P. Two of these mutations, D936Y and S943P, were among the most frequent in the ensemble of mutations we identified. Besides the widespread D614G, now dominant over the original D614 variant (37,38), only 5 other mutations (two of them being very peripheral, L5F and P1263L) recurred indeed in ≥ 20 sequences (see Table S1). S943P was also reported in (38), where it was hypothesized to be spreading via recombination.

The D936Y mutation was found in 25 sequences overall. In 22 sequences it was associated to the D614G mutation, while in 2 sequences it was associated to both the D614G and the A1020V mutations. The first D936Y/D614G variant was reported as a single sequence in USA (Washington) on March 15^th^, then, starting from March 17^th^ in England (7 sequences, March 17^th^ to 31^st^), Wales (7 sequences, March 17^th^ to 30^th^), the Netherlands (4 sequences, March 18^th^ to 29^th^) and Iceland (3 sequences, March 19^th^ to 28^th^). The 2 D936Y/D614G/A1020V variants were both reported from Wales, on March 26^th^ and 30^th^, therefore it might be hypothesized that they originated from the D936Y/D614G variant, already circulating in Wales at the time. In addition, the D936Y mutation was found in a unique S protein sequence, from France, dated March 18^th^, where it was not associated to the D614G variant.

The 22 sequences featuring the S943P mutation on April 21^st^ were all from Belgium. Twenty of them were associated to the D614G mutation and were reported between March 1^st^ and March 20^th^. In addition, two unique sequences featured the S943P mutation.

The S939F mutation was found associated to the D614G mutation in 8 sequences. It was first reported in 1 sequence from France on March 4^th^, then in 1 sequence from Iceland on March 16^th^, then again in 5 sequences from USA (Utah) between March 19^th^ and March 29^th^, then, finally, in 1 sequence from the Netherlands on April 2^nd^. In addition, this mutation was found in a unique S protein sequence, from Switzerland, dated February 26^th^, where it was not associated to the D614G variant.

The L938F mutation was a particularly late one; it was found in 2 sequences, associated to the D614G mutation, both from England and dated March 29^th^.

The S929I mutation was found in 2 sequences from USA (Washington), dated March 12^th^ and 27^th^, associated to the D614G mutation.

Finally, the S940F mutation had a unique geographical distribution, as it was found in 2 sequences from Australia (New South Wales) dated February 28^th^ and March 4^th^, not associated to the D614G mutation. In addition, it was found in 1 single sequence from France, dated March 20^th^, where it was associated to the D614G mutation.

In conclusion, with the exception of S940F, which was found in Australia, all the mutations in the HR1 core fusion were spread in two continents, Europe and/or North America.

Furthermore, most of them originated from the D614G variant. This is in agreement with them seeming to be quite late mutations, sequenced starting from the end of February/March 2020, i.e. over two months after the first Wuhan variant dated December 24^th^ 2019 (30) (Table 1). The frequency and geographical distribution of D936Y especially are noteworthy, considering that it was first sequenced only on March 15^th^.

**Table 1.** Occurrences of mutations on the HR1 “fusion core”.

|  | # Genomic/S protein<br>sequences | S929I | D936Y | L938F | S939F | S940F | S943P |
| --- | --- | --- | --- | --- | --- | --- | --- |
| **April 21<sup>st</sup>** | 10,823<sup>a</sup>/7,692<sup>b</sup> | 2 | 25 | 2 | 9 | 3 | 22 |
| **May 29<sup>th</sup>** | 34,805/23,332 | 5<sup>c</sup> | 213 | 3 | 20 | 5 | 3<sup>d</sup> |
| **% delta** | 233/203 | 150 | 752 | 50 | 122 | 67 | -633 |

<sup>a</sup> As downloaded from GISAID.
<sup>b</sup> Complete S proteins we extracted and translated (no undefined nucleotide, no internal stop codon, no insertion/deletion).
<sup>c</sup> In 2 additional sequences it is mutated to T.
<sup>d</sup> In 3 additional sequences it is mutated to I.

### Update on May 29^th^

On May 29^th^ 2020, we analysed the frequency of the above mutations and the emergence of other possible mutations in the HR1 fusion core in the updated GISAID dataset, containing 34,805 genomic sequences, from which we extracted 23,332 complete S protein sequences, thus roughly tripling the originally analysed April 21^st^ dataset. The result was someway surprising.

A positive selection pressure seemed to emerge for the D936Y mutation, which passed from the 25 cases of the April 21^st^ dataset to the 213 cases of the May 29^th^ dataset, corresponding to a ≈9-fold increment. Of the novel occurrences of the D936Y mutation, only 2 were found in USA (Utah and Minnesota), while most of them came from Europe, especially from UK, 68 from England, 56 from Wales and 1 from Scotland, and from Sweden, 55. Notably, the total number of occurrences of the D936Y mutation amounted to the 20.5% of all the 273 sequences available from Sweden, and to the 2.7% and 1.4% of all the sequences available from Wales and England (5397 and 2374, respectively). The remaining ones came from Denmark, 5, and Poland, 1.

The ≈3-fold increment in frequency of the S929I and S939F mutants was in line with the increment of the sequences in the dataset. The three additional occurrences of the S929I mutation were from USA (Washington), Wales and England. A novel S929T mutation was also reported twice from Scotland. Additional occurrences of the S939F mutation were instead from USA, 7, England, 2, Kazakhstan, 1, and UAE, 1. As for the L938F and S940F mutants, their increment was significantly lower than the increment of the sequences in the dataset. A positive selection thus clearly hasn’t emerged to date for these mutations. The only additional occurrence of L938F was from Denmark, while the 2 additional occurrences of S940F were from France and USA (Washington).

The S943P mutation represented a special case. Most of the sequences harbouring such a mutation were indeed modified between the April 21^st^ and the May 29^th^ datasets, so that they do not feature anymore the mutation to proline. However, 3 novel occurrences of the same mutation, S943P, emerged from China (Beijing). In addition, 3 sequences from Scotland presented the novel S943I mutation.

As for the remaining positions of the HR1 fusion core, to May 29^th^, either they were fully conserved (S937, K933, A942, A944, L945, G946, K947 and Q949), or they hosted one single occurrence of mutation (to valine for A930, to aspartate for I931 and G932, to histidine for Q934 and D935, to alanine for T941 and to arginine for L948). Because of the rarity of such mutations, we will not discuss them here. However, we will continue to monitor them over time.

### Sequence conservation

All the amino acids undergoing mutations in the SARS-CoV-2 S protein are conserved in the bat coronavirus RaTG13 S protein (sharing an overall sequence identity of 97% with SARS-CoV-2 S protein), while as many as five of them are mutated in the SARS-CoV-2 S protein (overall 76% sequence identical to the SARS-CoV-2 homolog) (see Figure 1). Four of these mutations are however conservative (aspartate to glutamate, serine to threonine), except S929, which is a lysine in SARS-CoV. It has been proposed that such mutations in the SARS-CoV-2 HR1 may be associated with enhanced interactions with the HR2, further stabilizing the 6-HB structure and maybe leading to increased infectivity of the virus (29). It is noteworthy that no one of the point mutations we identified restored the corresponding SARS-CoV amino acid.

### Effect of the mutations on the protein pre-fusion conformation

In the pre-fusion conformation, all the mutated positions, but S943, are located on the second of four non-coaxial helical segments composing the HR1 (Figure 1). Four of them, S929, D936, S939 and S940, are exposed to the solvent (Table 2), and can be modelled as larger (mostly aromatic) residues without causing any structural strain (see Figure 2). These mutations are not expected to cause relevant changes in the prefusion structure, although they could have a destabilizing effect as a consequence of posing large aromatic residues in direct contact with the solvent instead of smaller apolar (leucine), polar (serine in 2 cases) or even charged (aspartate) residues. In addition, S940 involves its side-chain in a H-bond with the main-chain of D936, 4 residues upstream. The loss of this H-bond in the S940F mutant also points to a slight destabilization of the pre-fusion conformation. As for L938, it is buried in the prefusion conformation, pointing towards a three-stranded anti-parallel β-sheet made of the S711-P728 segment from the S1 subunit and of the Y1047-P1053 and G1059-A1078 segments from the S2 subunit, without directly contacting it (distances above 5 Å). It can also be modelled as a large phenylalanine without causing sterical strain. Upon mutation, it seems to optimize the hydrophobic interactions with the neighboring residues, especially I726 and A944.

**Table 2.** Solvent accessibility of mutated residues in the pre- and post-fusion conformations.

| Amino acid | Pre-fusion | Post-fusion |
| --- | --- | --- |
| I929 | exposed | partly buried (18.6%)<sup>a</sup> |
| Y936 | exposed | partly buried (19.0%) |
| F938 | buried (95.8%) | buried (95.3 %) |
| F939 | exposed | exposed |
| F940 | exposed | buried (62.3 %) |

<sup>a</sup> Percentage of buried surface upon complex formation.

**Figure 2.**
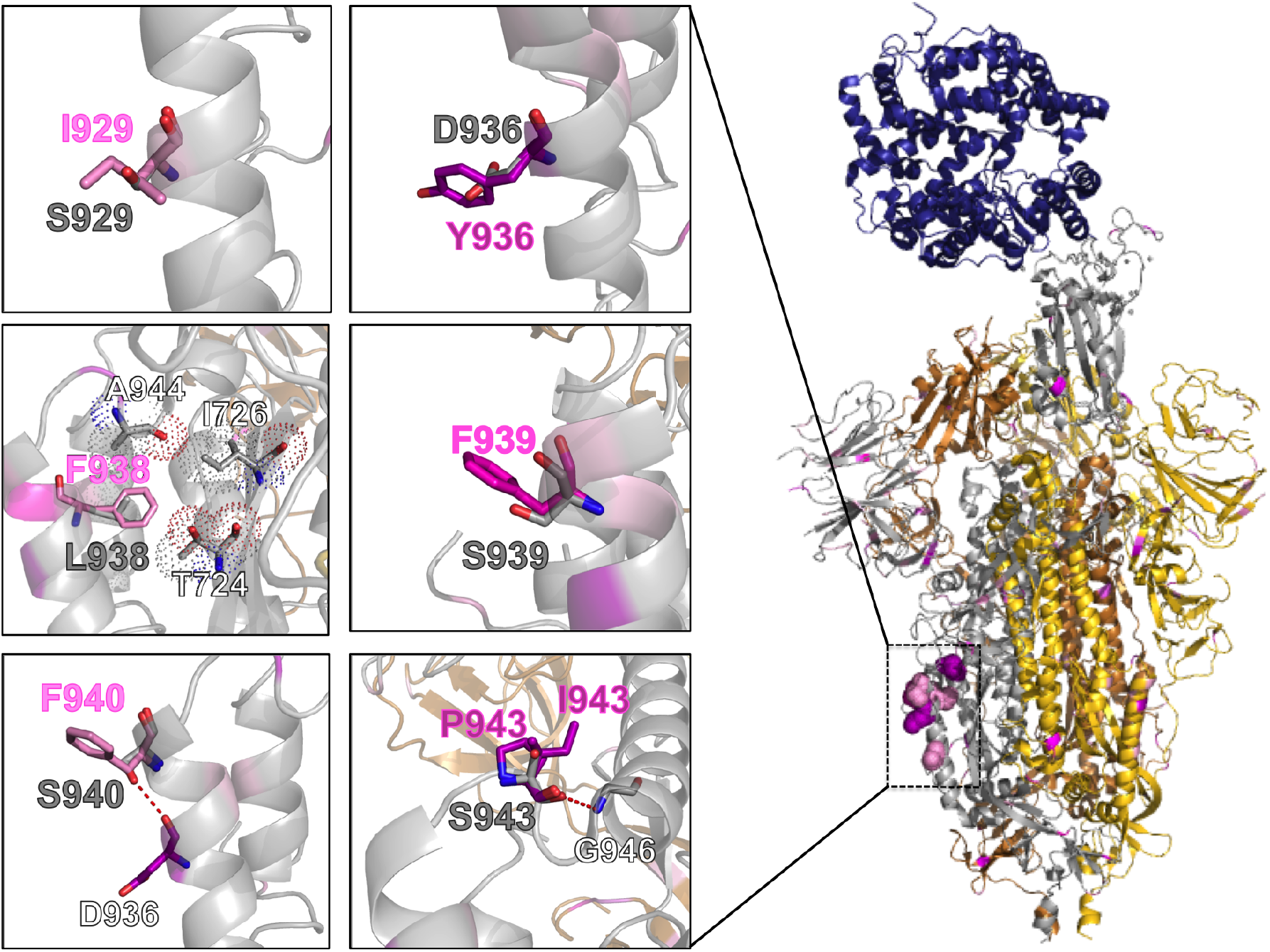
Models of mutants in the pre-fusion conformation. *Right:* Cartoon representation of the SARS-CoV-2 S protein in its pre-fusion trimeric conformation (the three monomers are colored in silver, gold and copper; PDB ID: 6VSB), with the structure of the RBD bound to the ACE2 receptor (in blue; PDB ID: 6M0J) superimposed on its chain A. All the mutated positions we identified in GISAID on April 21^st^ are colored on a purple-to-pink scale, depending on their relative frequency. Mutations in the HR1 fusion core are shown in a dots representation for chain A. *Left:* Focus on the structural context of each wild-type residue (silver sticks) and corresponding mutant (purple-to-pink sticks). Contacting residues (within 5 Å) are shown in a dots representation and H-bonds are shown as red dashed lines.

Finally, S943 is located on a turn immediately downstream the helical segment hosting the above five mutations, between the second and third helical segments. The wild-type residue S943 features ϕ and ψ dihedral angles of 58.5° and 24.5°, respectively, which fall in an unfavourable region for prolines. In the S943P model we generated, the P943 ϕ and ψ dihedrals assume the values of 3.0° and 68.2°, placing the residue in an outlier region (39). The favoured ϕ angle for prolines is indeed restricted to the value of −63 ± 15°, (40) characteristic of α-helices. A proline at such a position would therefore introduce an anomaly in the pre-fusion conformation, while strongly promoting the transition to the post-fusion single continuous helical conformation. It is also worth noticing that this would be the only mutation among those we identified so far, introducing a proline residue in the SARS-CoV-2 S protein (Table S1). In light of the analysis of the GISAID May 29^th^ updated, we also modelled the S943I mutation. Being isoleucine compatible with the S943 dihedral values, this mutation does not result in any structural strain.

### Effect of the mutations on the protein post-fusion conformation

When looking at the post-fusion conformation of the SARS-CoV-2 spike protein S2 subunit, these mutations appear more revealing. Three of the wild-type residues, S929, D936 and S943, are indeed engaged in side-chain to side-chain H-bonds with the HR2 segment of an adjacent monomer. In particular, S929, D936 and S943 (HR1 on Chain A) are H-bonded to S1196, R1185 and E1182, respectively (HR2 on Chain C, Figure 3). These are all strong H-bonds, especially the one between S943 and E1182, involving a negatively charged residue, and the one between D936 and R1185, being actually a salt bridge (estimated to contribute an additional 3-5 kcal/mol to the free energy of protein stability as compare to a neutral H-bond (41)). All these three H-bonds are lost upon mutation, which points to a weakening of the post-fusion assembly.

**Figure 3.**
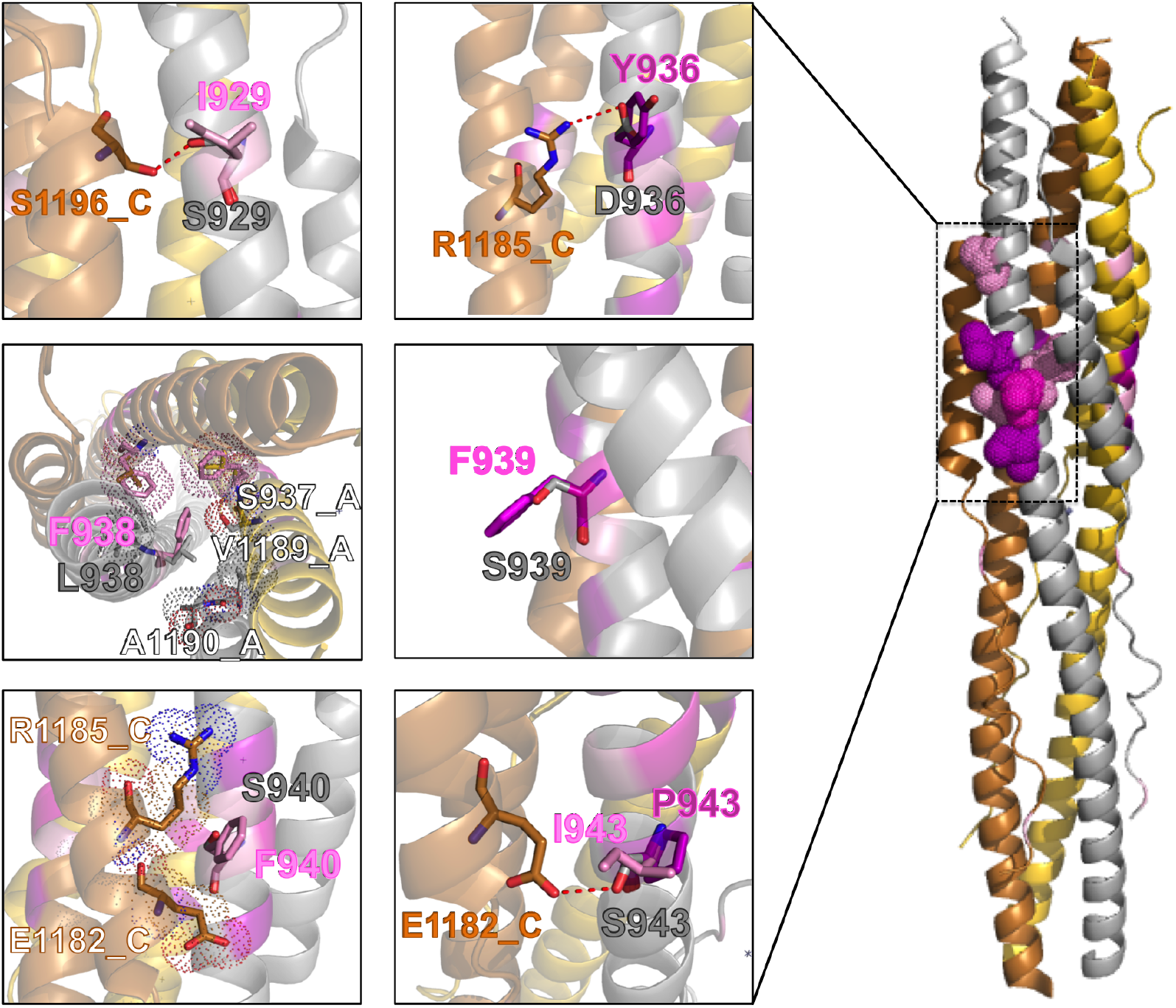
Models of mutants in the post-fusion conformation. *Right:* Cartoon representation of the SARS-CoV-2 S protein in its post-fusion trimeric conformation (the three monomers are colored in silver, gold and copper; PDB ID: 6LXT). The color code is the same of Figure 2. Mutations in the HR1 fusion core are shown in a dots representation for chain A. *Left:* Focus on the structural context of each wildtype residue (silver sticks) and corresponding mutant (purple-to-pink sticks). Contacting residues (within 5 Å) are shown in a dots representation and H-bonds are shown as red dashed lines.

Of the remaining three mutations, S939F is completely exposed to the solvent and therefore, like in the pre-fusion conformation, expected to act unfavourable on the protein solvation energy. On the contrary, in case of L938F and S940F, which are substantially buried within the structure, mutation to a large aromatic phenylalanine seems even to optimize the network of the hydrophobic interactions; in case of F940, with the aliphatic parts of the side-chains of E1182 and R1185 on an adjacent monomer, and, in case of F938, with V1189 and A1190 on the same monomer and with other F938 residues on both the adjacent monomers.

When comparing the effect of the mutations on the pre- and post-fusion structures, it emerges that the S929I, D936Y and S943I mutations strongly destabilize the postfusion conformation, while having a marginal impact on the stability of the pre-fusion one. On the contrary, S940F seems to favour the post-fusion conformation over the pre-fusion one. As for S938F and S939F, they seem to have a comparable effect on both the conformations, slightly stabilizing and destabilizing, respectively. Finally, the S943P mutation would strongly destabilize both the pre- and post-fusion conformations.

## Conclusions

Based on a thorough analysis of the S protein sequences, that we extracted from the genomic sequences of SARS-CoV-2 reported in GISAID on April 21^st^, we identified the fusion core of the HR1 as a mutational hotspot. The D936Y and S943P mutations were the most numerous, being among the most frequently occurring mutations overall at the time. Other, less frequent, mutations were S939F and then S929I, L938F and S940F. Overall, such mutations appeared to be late ones, emerging starting from the end of February or even mid March 2020, and were mainly localized in Europe and USA. Based on their frequency, on their location in a protein region playing a key role in the post-fusion conformation and also on the non-conservative nature of the mutations themselves, we decided to further investigate the structural basis of such mutations, finding out that they all can play a role in tuning the stability of the pre- and/or post-fusion S protein conformation.

A search of the GISAID dataset updated to May 29^th^ revealed a ≈9-fold increase of the D936Y mutation over time (*versus* a ≈3-fold increase in the dataset sequences of 203%), thus indicating a possible positive selection for it. Notably, the D936Y variant represented the 20.5% of all the sequences reported from Sweden and the 2.7 and 1.4 % of all the sequences form Wales and England, respectively.

Other potentially interesting mutations are S929I and S939F, whose number of occurrences underwent a ≈2/3-fold increase. On the other hand, the increment in the occurrence of L938F and S940F was marginal, posing less emphasis on such mutations, which will be nonetheless useful to continue monitoring.

Finally, the S943P mutation, although still reported in few cases, underwent a dramatic reduction of occurrences, due to modification of the original sequences where they were first reported. At the same time, a S943I mutation emerged, that will also be worth continuing to monitor. We remind here that a proline at position 943 would cause a significant destabilization on the S protein pre-fusion conformation.

It is also worth noticing that the 2 mutations significantly increasing their frequency over time, D936Y and S929I, were also those that, together with S943P/I, caused the loss of a inter-monomer H-bond in the post-fusion conformation of the protein. Interestingly, the now emerging S943I mutation gets the same effect without destabilizing the pre-fusion conformation. The most frequently occurring mutation in the HR1 “fusion core”, common in Sweden and UK on May 29^th^, is also the one causing the loss of a strong inter-monomer salt bridge. Our structural analyses provide a rationale for such mutations, pointing to a weakening of the post-fusion assembly. However, only experiments on cellular systems will clarify whether this may be a virus strategy for reducing its membrane fusion capacity, thus lowering its virulence.

## Acknowledgements

We gratefully acknowledge all the Authors from the Originating laboratories responsible for obtaining the specimens and the Submitting laboratories where genetic sequence data were generated and shared via the GISAID Initiative, on which this research is based.

R.O. thanks MIUR-FFABR (Fondo per il Finanziamento Attività Base di Ricerca) for funding; L.C. acknowledge King Abdullah University of Science and Technology (KAUST) for support and the KAUST Supercomputing Laboratory for providing computational resources.

## Authors’ contribution

L.C. participated in the study’s design and carried out the analyses. R.O. conceived of the study, participated in its design, carried out the analyses and drafted the manuscript. All authors read and approved the final manuscript.

---

**Table S1.**
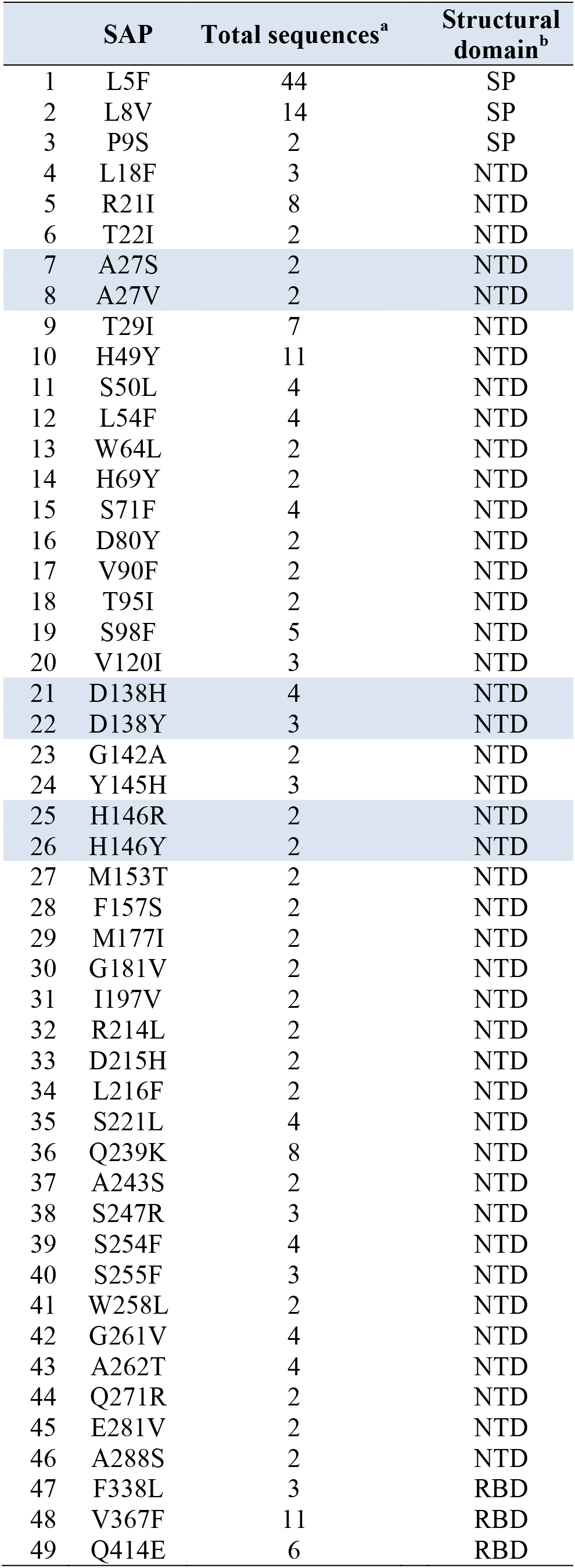
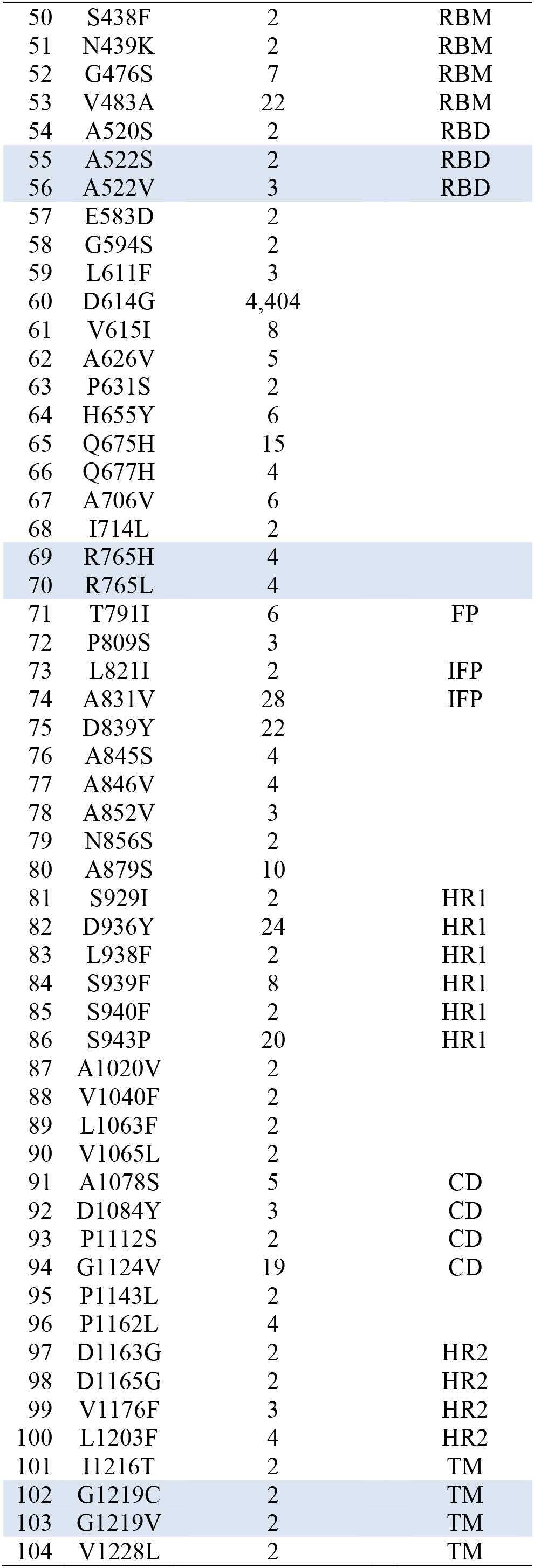
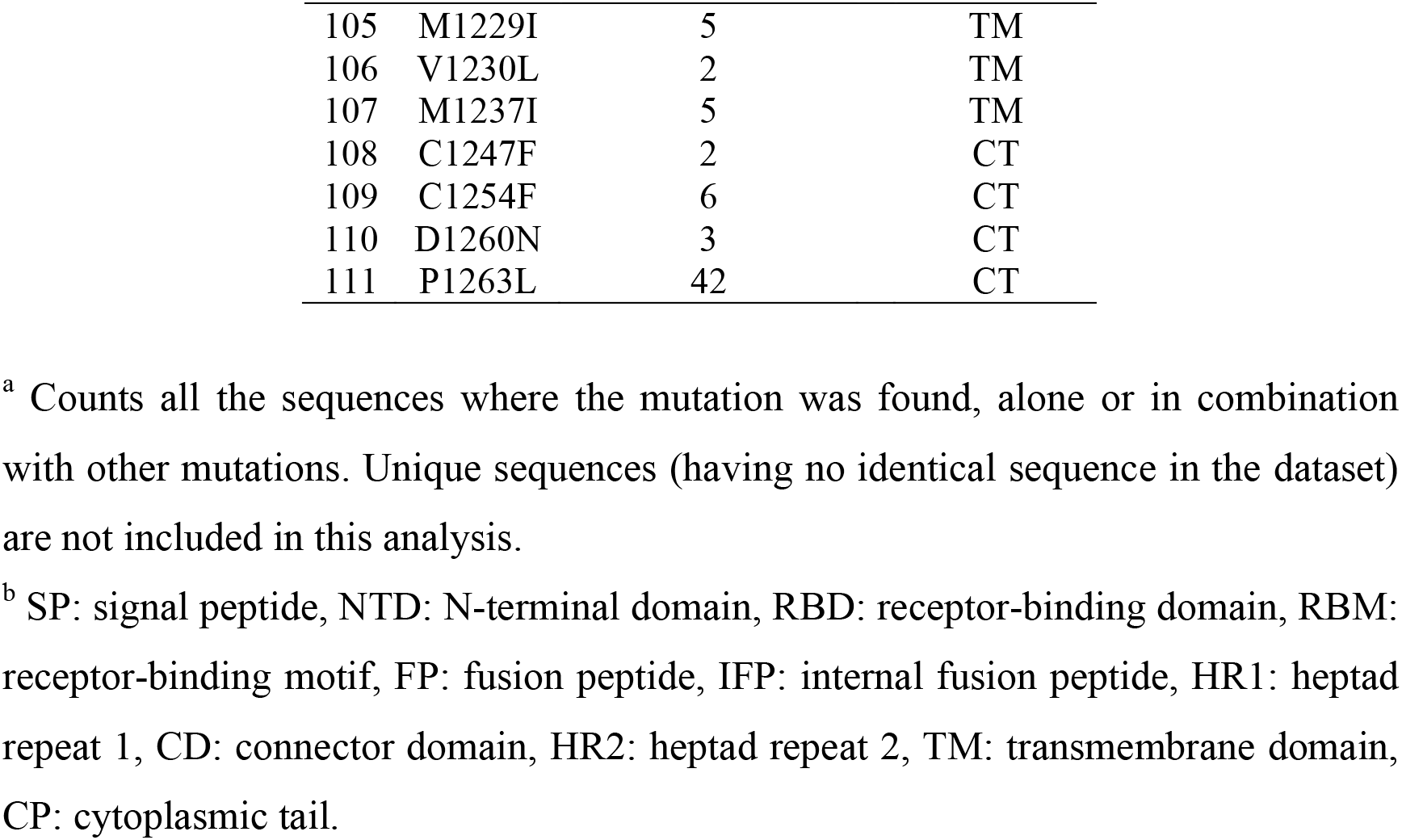
List of mutations identified in GISAID in at least 2 identical sequences on April 21^st^ 2020, in sequential order.

